# A regulatory domain in the K_2P_2.1 (TREK-1) carboxyl-terminal allows for channel activation by monoterpenes

**DOI:** 10.1101/2020.01.14.906396

**Authors:** Eden Arazi, Galit Blecher, Noam Zilberberg

## Abstract

Potassium K_2P_ (‘leak’) channels conduct current across the entire physiological voltage range and carry leak or ‘background’ currents that are, in part, time- and voltage-independent. K_2P_2.1 channels (i.e., TREK-1, KCNK2) are highly expressed in excitable tissues, where they play a key role in the cellular mechanisms of neuroprotection, anesthesia, pain perception, and depression. Here, we report for the first time that human K_2P_2.1 channel activity is regulated by monoterpenes (MTs). We found that cyclic, aromatic monoterpenes containing a phenol moiety, such as carvacrol, thymol and 4-IPP had the most profound effect on current flowing through the channel (up to a 6-fold increase). By performing sequential truncation of the carboxyl-terminal domain of the channel and testing the activity of several channel regulators, we identified two distinct regulatory domains within this portion of the protein. One domain, as previously reported, was needed for regulation by arachidonic acid, anionic phospholipids and temperature changes. Within a second domain, a triple arginine residue motif (R344-346), an apparent PIP_2_-binding site, was found to be essential for regulation by holding potential changes and important for regulation by monoterpenes.

## 1 Introduction

Potassium channels selectively and rapidly enable the movement of K^+^ ions across biological membranes down the electrochemical K^+^ gradient at a rate close to the rate of diffusion (MacKinnon, 2003). Members of the potassium leak channel family are structurally unique among potassium channels as each subunit possesses four transmembrane segments and two pore-forming domains (2P/4TM). As such, these channels are often referred to as two pore-domain K+ or K_2P_ channels (Choe, 2002; Goldstein et al., 2001). Since the cloning of the *Drosophila melanogaster* K_2P_ channel KCNK0(Goldstein et al., 1996), genes encoding putative potassium leak subunits have been recognized in sequence databases of all eukaryotes (Lesage et al., 2000), including 15 genes in the human genome (Talley et al., 2003).

K_2P_ channels conduct current across the entire physiological voltage range and carry leak or ‘background’ currents that are, in part, time- and voltage-independent. These channels are essential to neurophysiological function as their activity suppresses excitability, in part, through the maintenance of a resting membrane potential below the threshold for action potential firing (Goldstein et al., 2001). Hence, K^+^ leak currents shape the duration, frequency and amplitude of action potentials, and, therefore, modulate cell responsiveness and excitability (Goldstein et al., 2001; Hille, 2001; Hodgkin and Huxley, 1952). It was shown that K_2P_ channels can also increase excitability by supporting high-frequency firing once an action potential threshold is reached (Brickley et al., 2007). It was recently reported that the majority of K_2P_ channels are gated by membrane potential, in spite of their lack of a voltage sensor, as the outward current of K^+^ ions through the selectivity filter was found to open this gate (Schewe et al., 2016).

K_2P_ channels have been shown to participate in and modulate various important physiological processes, such as pain perception (Alloui et al., 2006) and cardiac activity (Decher et al., 2017; Schmidt et al., 2012; Schmidt et al., 2017). The human K_2P_2.1 (TREK-1, KCNK2) channel is expressed at high levels in excitable tissues, such as the nervous system (Fink et al., 1996), heart(Aimond et al., 2000), and smooth muscle (Koh et al., 2001). K_2P_2.1 channels function as signal integrators, responding to a wide range of physiological and pathological inputs, as their activity and biophysical properties are strongly modulated by various physical and chemical signals (Bagriantsev et al., 2012; Cohen et al., 2008; Cohen et al., 2009; Honore, 2007; Noel et al., 2011). K_2P_2.1 was also shown to be involved in numerous physiological processes, such as pain perception (Alloui et al., 2006; Cohen et al., 2009; Li and Toyoda, 2015), general anesthesia and neuroprotection (Franks and Honore, 2004). In addition, K_2P_2.1 was found to be a target of fluoxetine (Heurteaux et al., 2006) and its activity was enhanced by mood stabilizers, such as lithium-chloride, gabapentin, valproate, and carbamazepine (Kim et al., 2017). The elective opening of this channel led to a reduction in dorsal root ganglion (DRG) neuron excitability (Loucif et al., 2017). K_2P_2.1 is strongly activated by neuroprotective agents, such as arachidonic acid and riluzole (Noel et al., 2011). In the heart, deficiency of K_2P_2.1 led to defects in sinoatrial cell membrane excitability (Unudurthi et al., 2016). Finally, K_2P_2.1 was found to play a role in the perception of a variety of pain stimuli, including thermal and mechanical pain, and co-localized with the thermal sensor TRPV1 in small DRG neurons (Alloui et al., 2006; Pereira et al., 2014).

Terpenes are a large group of organic chemicals that are mostly produced in plants. The basic building block of terpenes and terpenoids (modified terpenes) is the five-carbon moiety isoprene, with the number of isoprene units used, serves to classify these molecules. Linear or cyclic monoterpenes (MTs) consist of two isoprene units. MTs have been known for centuries for their beneficial effects as anti-fungal (Marei et al., 2012), anti-bacterial (Garcia et al., 2008) and analgesic (Khalilzadeh et al., 2016) agents. Terpenes have been long recognized as remedies for pain (Guimaraes et al., 2014; Quintans-Junior et al., 2013; Quintans Jde et al., 2013) and cardiovascular diseases (Aydin et al., 2007; Magyar et al., 2004; Menezes et al., 2010; Peixoto-Neves et al., 2010; Santos et al., 2011), as well as for their anti-tumor, local anesthetic and anti-ischemic abilities (Koziol et al., 2014). Research into therapeutic uses for the terpenes found in cannabis and terpenophenolic cannabinoids has increased over the last three decades (Owens, 2015). Since the essential oils content in plants is up to 3.5% of their dry weight and as MTs comprise up to 40% of the essential oils (Al-Kalaldeh et al., 2010; Arslan, 2016), the MTs consumption in a normal diet is rather low. High concentrations of MTs in plasma and certain tissues were reported in chickens fed with high amounts of MTs (Ocel’ova et al., 2016), and in rats, intravenously injected or fed with an emulsified MT (Pavan et al., 2018).

Several MTs were found to have an effect on ion channels, both in excitable cells (Oz et al., 2015) and in other tissues (Murbartian et al., 2005). Carvacrol (2-methyl-5-(propan-2-yl) phenol), found in oregano, and thymol, found in thyme, activates and sensitizes the murine and human transient receptor potential (TRP) vanilloid TRPV3 channel. Acyclic MTs, like citronellol, nerol, and their derivatives, were found to modulate the activity of TRPA1 (Ortar et al., 2014). (-)-menthol, (+)-menthol, as well as cineole, carvacrol, and thymol, reduced compound action potential peaks in frog sciatic nerve by inhibition of voltage-gated ion channels, like TTX-sensitive sodium channels, or by non-specific interaction with the lipid membrane (Kawasaki et al., 2013). α-Thujone was found to inhibit almost all GABA receptor types (Czyzewska and Mozrzymas, 2013). Carvacrol inhibits the mammalian TRPM7 channel (Parnas et al., 2009) while activating the human TRPV3 and rat TRPA1 channels (Haoxing Xu, 2006). Eugenol activates the murine TRPV3 but not the TRPV2 or TRPV4 channels (Haoxing Xu, 2006). To date, the response of any of the K_2P_ channels to MTs has not been explored. Since terpenes are known to reduce pain sensation and since activation of K_2P_2.1 channels was shown to reduce pain, we examined whether K_2P_2.1 channels are activated by any of the MTs.

Here, we demonstrate, for the first time, that MTs are powerful modulators of the human K_2P_2.1 channel. Furthermore, we located a four amino acid-long domain in the carboxyl-terminal domain that is required for channel activation. We also discuss the structural and physical properties of MTs that contribute to their ability to influence this channel.

## 2. Methods

### 2.1. Animals

All experiments using animals were performed in accordance with the guidelines of the institutional animal care and use committee. The project approval number is IL-61-09-2015.

### 2.1. Cloning

Channels were cloned into plasmid pRAT that includes a T7 RNA polymerase promoter to enable cRNA synthesis, as well as the 3’-UTR and 5’-UTR sequences of the *Xenopus laevis* β-actin gene to ensure efficient expression in *Xenopus* oocytes. Competent *Escherichia coli* DH5α cells were transformed by heat shock. Plasmid DNA was purified with a Wizard Plus SV Miniprep kit (Promega). Restriction enzyme digestions were performed according to the manufacturer’s instructions (Thermo Fisher Scientific or NEB). Point mutations were generated according to the Quickchange site-directed mutagenesis technique (Stratagene) and confirmed by sequencing. PCR amplifications were performed using a PTC-2000 instrument (MJ Research) with PFU DNA polymerase (Thermo Fisher Scientific). cRNA was transcribed *in vitro* by T7 polymerase using mMESSAGE mMACHINE (Ambion) or the AmpliCap High Yield Message Maker (Epicentre) kits.

### 2.2. Electrophysiology

*Xenopus laevis* oocytes were isolated and injected with 20-40 nL of solutions containing 0.3–40 ng cRNA using a 3.5’’ Drammond#3-000-203-G/X glass capillary, pulled in a Sutter P97 capillary puller, and a Drummond manual oocyte microinjection pipette (3-000-510-X). Whole-cell currents were measured 1– 3 days after injection by the two-electrode voltage-clamp technique (GeneClamp 500B, Axon Instruments). Data were filtered at 2 kHz and sampled at 5 kHz with Clampex 9.0 software (Axon Instruments). For two-electrode voltage-clamp experiments, the pipette contained 3 M KCl, and the bath solution contained (in mM) unless otherwise noted: 4 KCl, 96 NaCl, 1 MgCl_2_, 0.3 CaCl_2_, 5 HEPES, pH 7.4, with NaOH (standard solution). If required, the standard bath solution was supplemented with the solvent (ethanol) of the tested chemical.

Injection of cRNA into oocytes was done in OR-2 solution (in mM: 5 HEPES, 1 MgCl_2_, 2.5 KCl, 82.5 NaCl, pH=7.4). Post-injection oocytes were maintained in ND-91 solution (in mM: 5 HEPES, 1 MgCl_2_, 1.8 CaCl_2_, 2 KCl, 91 NaCl, pH=7.4). Injection of carvacrol and isopropylphenol into oocytes during electrophysiological recording was done as follows: An injection solution (up to 75 mM) in OR-2 solution was prepared from either carvacrol or isopropylphenol stock solutions. A glass capillary, prepared as described above, was filled with the injection solutions. For injection, oocytes with an approximate volume of 1 microliter were selected. The injection was performed during voltage clamp, when 20 nl of each solution were injected to a final concentration of up to 3mM. Injections were administered gradually over 2-5 seconds. As a control for injection quality, 20 nl of 125 mM BaCl_2_ in OR-2 solution were injected into control oocytes, resulting in a non-specific rise in outward current (not shown). To determine the voltage-dependent fraction of the current, the initial, voltage-independent (instantaneous) current was estimated by fitting the current to an exponential decay slope as the initial currents are masked by the capacitive transient current.

### 2.3. Chemicals

Carvacrol (cat#282197), thymol (cat#T0501), p-cymene (cat#C121452), 4-isopropylphenol (cat# 175404), eugenol (cat#E51791, cinnamaldehyde (cat#W228613), menthol (cat#M2772), beta-citronelol (cat# C83201), geraniol (cat#16333, 4-methylcatechole (cat# M34200) and arachidonic acid (cat#A3611) were all purchased from Sigma-Aldrich.

#### 2.3.1. Preparation of compounds

Compounds delivered as powders were dissolved into stock solutions (4-6 M) in 100% ethanol. Compounds delivered as liquid oils (6.5-7.5 M) were diluted 1:1 with ethanol to form stock solutions. Stock solutions were kept at −20°C for up to two weeks. Just prior testing, stock solutions were diluted in the bath solution to the desired concentration. Diluted compounds were vigorously vortexed until completely dissolved. All solutions were supplemented with ethanol to a final concentration of 0.1% (v/v) (confirmed not to harm the oocytes). The pH was corrected to 7.4±0.05 using NaOH or HCl.

### 2.4. Statistical analysis

Data were expressed as the mean ± standard error of the mean (SEM) and analyzed and presented using Microsoft Excel 2016. Groups of two were analyzed using Student’s t-test. Values were considered to be significantly different when the P-value was less than 0.05. All experiments were repeated with at least five oocytes.

## 3. Results

### 3.1. Carvacrol is a novel K_2P_2.1 activator

Carvacrol was found to reduce the activity of voltage-gated sodium channels, to show anti-nociceptive and analgesic properties in mice (Chiu et al., 2011; Guimaraes et al., 2012; Silva et al., 2016) and to block the generation of action potentials in intact rat DRG (Joca et al., 2012). In addition, carvacrol was shown to promote neuroprotection in several animal models (Li et al., 2015; Yu et al., 2012). K_2P_2.1 is a member of the K_2P_ family that modulates the sensation of pain and heat and which is expressed in high quantities in sensory neurons of the DRG (Alloui et al., 2006). Furthermore, K_2P_2.1 activation is associated with neuroprotection of human and murine neurons (Heurteaux et al., 2004; Vallee et al., 2012). Therefore, we considered the effect of carvacrol on human K_2P_2.1 (hK_2P_2.1) channels. When expressed in *Xenopus laevis* oocytes, carvacrol dramatically enhanced K_2P_2.1-generated current when applied externally (Fig. 1A-D), yet had no effect when injected into the oocyte (Fig. 1A). K_2P_2.1 channel activation by carvacrol was dose-dependent, with an effective concentration of 50% (EC_50_) of 0.20±0.06mM (Fig. 1B). Menthol similarly affected K_2P_2.1 channel-generated current, albeit with a higher EC_50_ value (Fig. 1B). Furthermore, externally added carvacrol changed the reversal potential of the channel (Fig. 1E) by 5.6±1.0 mV (n=8-11). Two types of analyses were performed. To avoid bias due to current level changes, we included in our calculations only oocytes with comparable current levels (5-10 µA) either before or after incubation with carvacrol (Fig. 1E). Subsequent, we analyzed paired results from the same oocyte, regardless of its initial or the final current, and obtained similar results (reversal potential change of 5.3±1.1 mV, n=17; not shown). In addition, carvacrol reduced the voltage-dependent current by 19.5±1.2% (Fig. 1F).

**Fig. 1.**
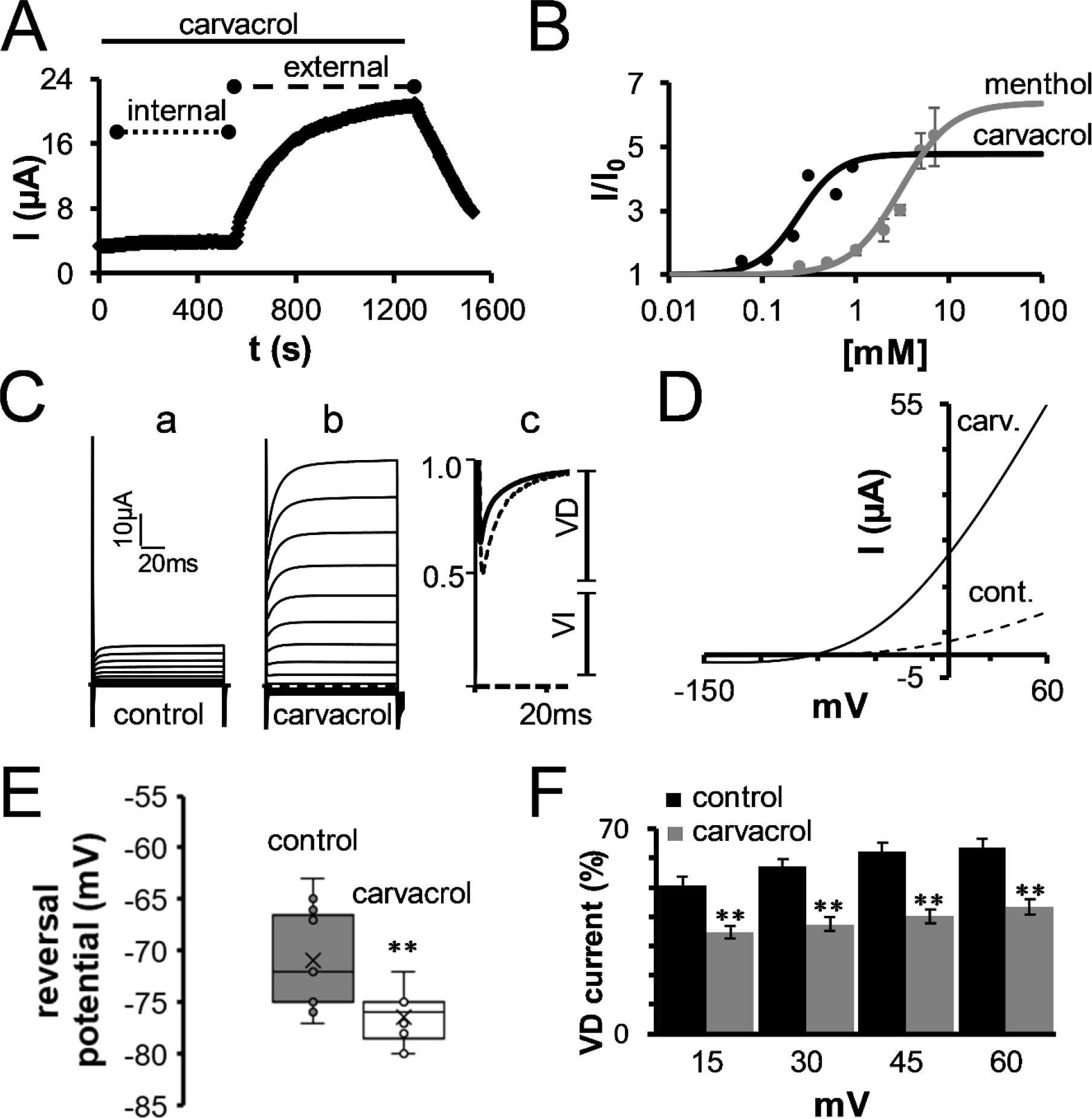
Activation of K_2P_2.1 by monoterpenes. (A) Only externally applied carvacrol activates K_2P_2.1 channels. Currents from a representative oocyte expressing K_2P_2.1 channels were measured by the TEVC technique before and after injection of carvacrol to a final internal concentration of 0.9 mM (internal) and during application of 0.3 mM carvacrol to the external side of the oocyte (external). Oocyte membrane potential was held at −80 mV and pulsed to +25 mV for 75 ms with 5 s interpulse intervals. Injection of carvacrol to a final concentration of up to 3mM resulted in similar results. (B) Concentration-dependence for groups of 5-10 oocytes, normalized to currents measured under control conditions (mean±SEM) for carvacrol (●) or menthol (●). Solid lines represent a fit of the data to the Hill equation. EC_50_ values of 0.20±0.06 mM and 3.8±1.0 mM were calculated for carvacrol and menthol, respectively. (C) Currents of a representative oocyte before (a) and after (b) application of 0.3 mM carvacrol. Oocyte membrane potential was held at −80 mV and pulsed from −150 to +60 mV in 15 mV voltage steps for 130 ms with 2 s interpulse intervals. (c) Normalized currents at 60mV of a representative oocyte displaying the voltage-dependent (VD) and the voltage-independent (VI) currents, before (dashed line) and after (solid line) application of carvacrol. (D) K_2P_2.1 steady-state current-voltage relationships measured as in C, before (dashed line) and after (solid line) application of 0.3 mM carvacrol. (E) Changes in reversal potential during the application of 0.3mM carvacrol. (F) Changes in voltage-dependent current (VD) at four holding potentials, during carvacrol application **P<0.01. VD current was calculated as the fraction of the time-dependent current from the total current.

### 3.2. Structural and physical properties of monoterpenes that determine their activity towards K_2P_2.1 channels

To determine whether other MTs activate K_2P_2.1 channels, we tested nine additional compounds with various chemical properties. The compounds chosen differed from one another in terms of their polarity, partition coefficient, and structure. Accordingly, we tested linear compounds (geraniol and β-citronellol), compounds with a benzene ring (carvacrol, eugenol, 4MC, p-cymene, 4-IPP and thymol) and compounds with a cyclohexane ring (menthol). The tested compounds also varied in terms of the number of hydroxyl groups. At a concentration of 0.3 mM, eight compounds were found to activate K_2P_2.1 channels (30-600% increased current; Fig. 2). We found that among the tested compounds, carvacrol was the most effective activator. Thymol, an isomer of carvacrol that differs only in the location of the hydroxyl group and 4-IPP that lacks one of the methyl groups activated the channel to a lesser extent. Linear compounds (geraniol and β-citronellol), compounds lacking a hydroxyl group (p-cymene), compounds lacking a benzene ring (menthol) and eugenol were much less active.

**Fig. 2.**
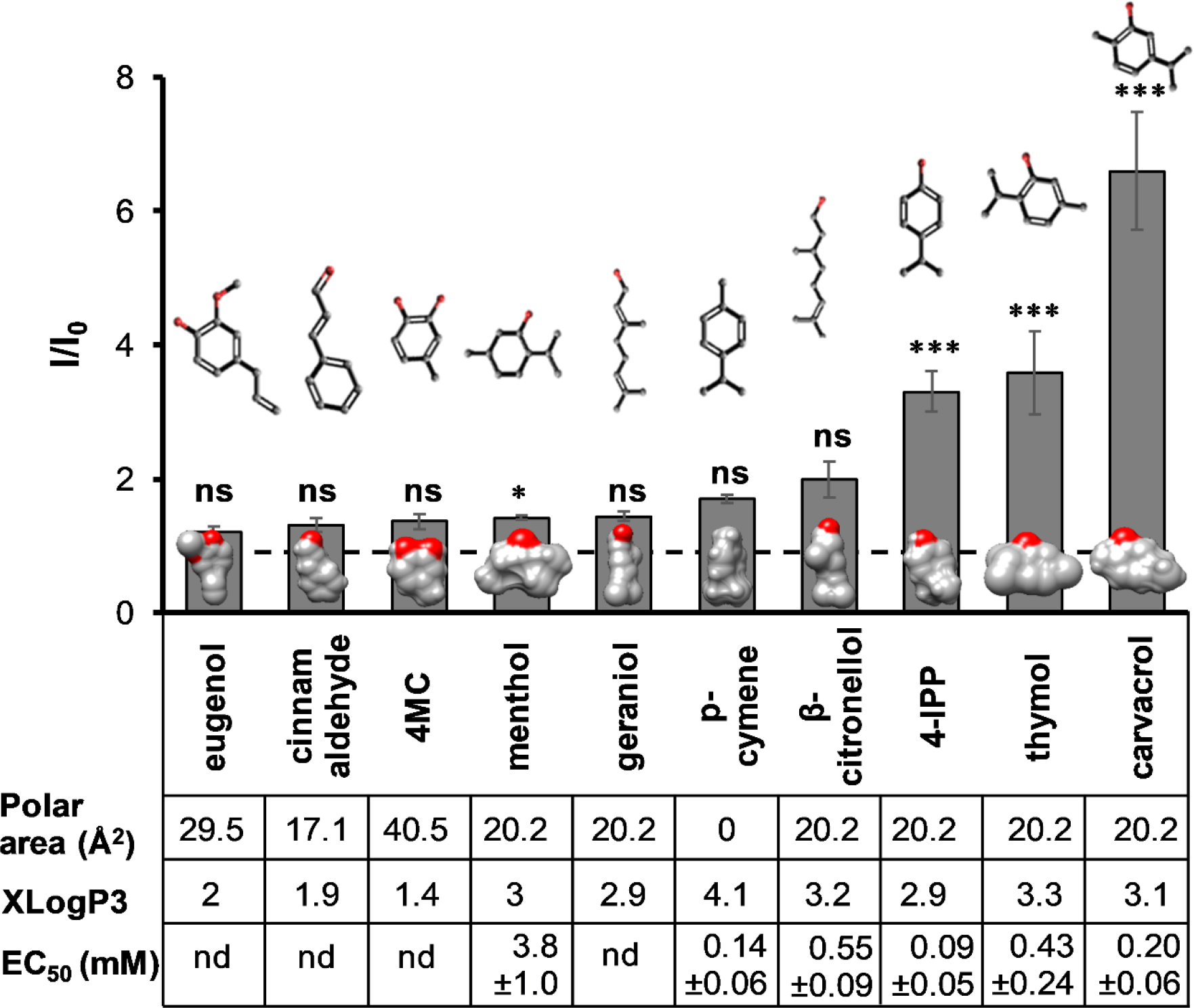
Chemical requirements for activation by monoterpenes. Activation of K_2P_2.1 channel currents was studied as in Fig. 1A (n=5-10, mean±SEM). All MTs were applied at 0.3 mM. Polar area (Å^2^), octanol-water partition coefficient (logP) prediction (XLogP3), 2D structures and the coordinates of the 3D structures of the terpenes were obtained from PubChem (Kim et al., 2019). 3D models were performed with the UCSF Chimera package (Pettersen et al., 2004). Oxygen molecules are colored red. The statistical significance of the difference from the initial current post compound application is denoted by asterisks. ns-not statistically significant. EC_50_ values were determined to compounds that at the concentration of 1mM increased the current by 50% and above. nd-not determined.

### 3.3. Activation of K_2P_2.1 by monoterpenes is independent of its phosphorylation state

Phosphorylation of the K_2P_2.1 channel carboxyl-terminal residue, Ser348 (Fig. 3A), by protein kinase A was found to be a major regulatory signal in this channel. Phosphorylation of Ser348 decreased the open probability (*P*o) of the channel (Maingret et al., 2000) and imposed a voltage-dependent phenotype onto the channel (Bockenhauer et al., 2001). To examine the importance of phosphorylation to channel activation by MTs, we tested two K_2P_2.1 mutants, K_2P_2.1-S348D and K_2P_2.1-S348A, mimicking the phosphorylated and the non-phosphorylated states, respectively. Both mutants were affected by carvacrol and menthol (Fig. 3B i), albeit to different extents. Since the mutations enforce the tendency of the channel to display either higher (A) or lower (D) basal activities (Bockenhauer et al., 2001), respectively, it is plausible that the inability of terpenes to open K_2P_2.1-S348A is mainly due to its initial high open probability. We thus looked at the maximal current levels of the wild type and the two mutant channels injected with the same amount of complementary RNA (cRNA) and currents were measured at the same approximate time (Fig. 3Bii). Since the final current level of all three channels was similar (student’s t-test, P>0.35), regardless of the initial current, we concluded that the phosphorylation state of K_2P_2.1 does not affect its responsiveness to terpenes.

**Fig. 3.**
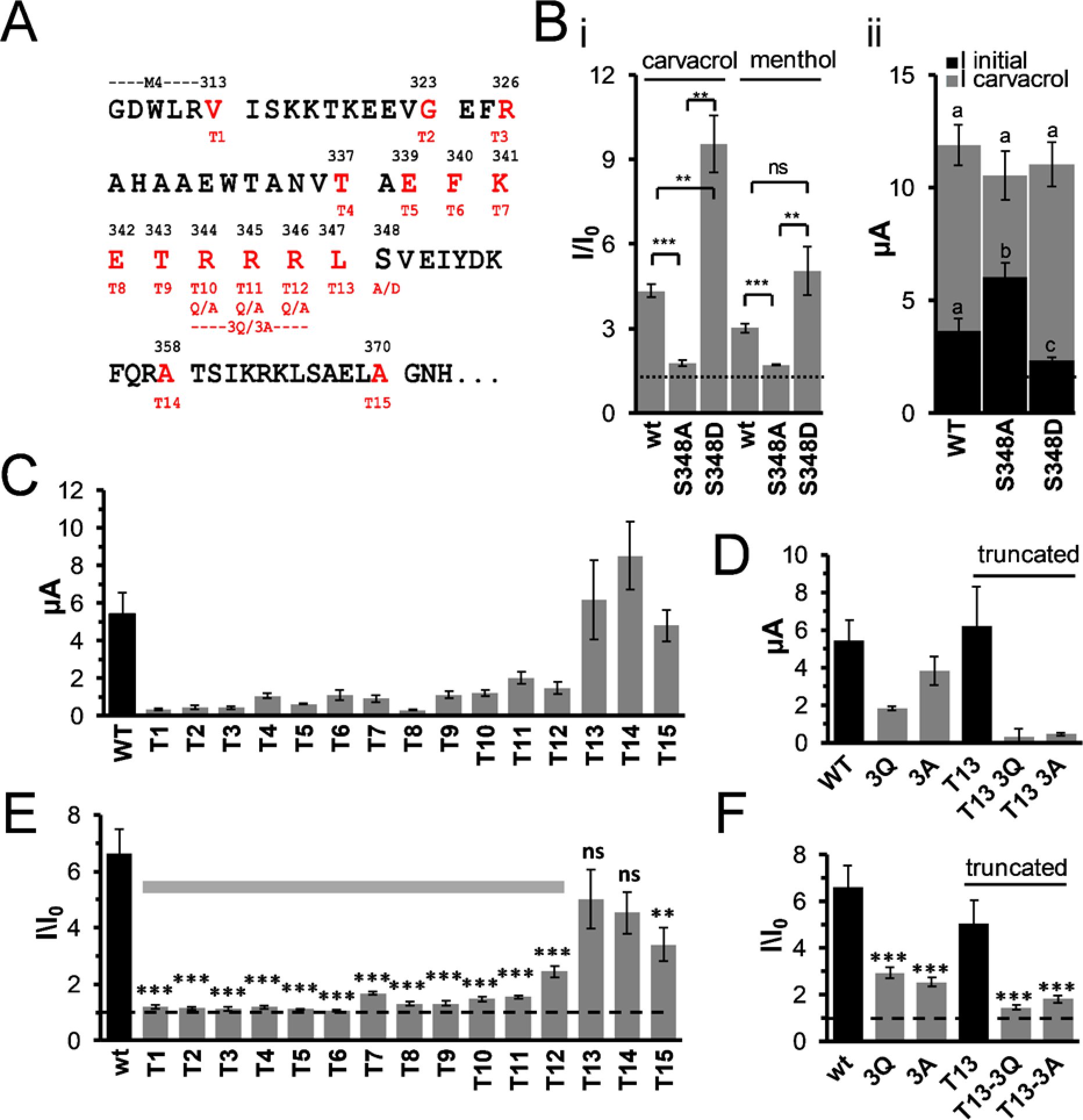
The C-terminal is essential for K_2P_2.1 monoterpenes activation. (A) Truncation (marked in red T and truncation number) and mutants (marked in the corresponding mutation) of the C-terminal domain of K_2P_2.1 channels and mutants. Mutated residues are in red. (B) (i) the current increase in wild-type K_2P_2.1 channels (wt), and in K_2P_2.1-S348A (S348A, “non-phosphorylated”), and K_2P_2.1-S348D (S348D, “phosphorylated”) mutant channels after application of 0.3 mM carvacrol or 3 mM menthol, as indicated. (ii) current before (I initial), and post-administration of 0.3 mM carvacrol (I carvacrol) (n=5-10, mean±SEM). As initial currents were significantly different (student’s t-test p value<0.05), while final currents were not (p>0.05), we concluded that the difference in current increase between the three channels was mainly due to the effect of phosphorylation on their basal currents and not on their responses to carvacrol. (C) Initial current levels of channel truncation mutants termed as in A (n=5-10, mean±SEM). (D) Initial current levels of full (3A/3Q) or truncated (T13 3A/3Q) channels mutated in the three-arginine cluster (n=5-10, mean±SEM). (E) Channel truncation mutants, termed as in A, were incubated with 0.3 mM carvacrol. The dashed line represents no change from the initial current. The horizontal bar indicates the minimal C-terminal domain that is essential for the response to carvacrol. Elongation of the C-terminal to 34 residues (T13) fully restored the response to carvacrol (student t-test p>0.05). (F) Replacing the three arginine cluster in full-length (3A/3Q) or truncated (T13 3A/3Q) channels significantly reduced the current in response to 0.3mM carvacrol (p<0.05).

### 3.4. The C-terminal domain of the K_2P_2.1 channel is essential for activation by carvacrol

In many members of the K_2P_ channel family, the cytoplasmically oriented C-terminal domain is essential for the proper regulation of channel activity. For example, while the activity of the *Drosophila* KCNK0 channel (K_2P_0) is enhanced by protein kinase phosphorylation (Zilberberg et al., 2000), removal of the C-terminal domain results in a channel that is insensitive to such activation (Cohen and Zilberberg, 2006; Zilberberg et al., 2000). The C-terminal domain of K_2P_2.1 channels was shown to be vital for regulation by phosphorylation (Bockenhauer et al., 2001; Maingret et al., 2000), temperature (Bagriantsev et al., 2012; Maingret et al., 2000), arachidonic acid (Patel et al., 1998), internal acidification (Honore et al., 2002), internal lysophosphatidic acid (Gonzalez et al., 2015), PIP_2_ levels (Chemin et al., 2005; Woo et al., 2018) and mechanical stretch (Maingret et al., 1999). It was suggested that the proximal part of the C-terminal domain, which contains a charged region, is critical for chemical and mechanical activation of the channel (Patel et al., 1998). This cluster of basic and acidic residues is central for channel modulation by negatively charged phospholipids and pH (Honore et al., 2002; Treptow and Klein, 2010). Two phosphorylation sites within this region were shown to regulate channel activation (Murbartian et al., 2005) via the addition of a negative charge to the otherwise positively charged region (Bockenhauer et al., 2001). It was later suggested (Chemin et al., 2005) and demonstrated (Sandoz et al., 2012) that various positive modulators of channel activity, such as internal pH, Gq-coupled receptors and phospholipids, act by tightening the interaction of the positively charged region with the inner leaflet of the plasma membrane. Recently, a triple arginine cluster was reported as being essential for the regulation of K_2P_2.1 by PIP_2_ (Woo et al., 2018).

We, therefore, addressed the role of the K_2P_2.1 C-terminal domain in the response of the channel to MTs. Complete deletion of the C-terminal domain abolished the response of K_2P_2.1 to carvacrol (T1 to T3 in Figures 3A, 3C, and 3E), as well as to thymol, beta citronellol and p-cymene (not shown), suggesting that the C-terminal domain is crucial for the activation of K_2P_2.1 by MTs.

### 3.5. Defining the region of the C-terminal domain that regulates activation by monoterpenes

To precisely define the region of the K_2P_2.1 carboxyl-terminal domain responsible for reacting to increased MT levels, an array of serial truncations was generated (Fig. 3A). All truncated channels, in the exception of T1, were functional, albeit with varying initial current levels (Fig 3C). We found that while the minimally functional channel (T2) was not affected by carvacrol (Fig. 3E), elongation by 23 amino acids (T12) partially restored the response (Fig. 3E), while the addition of one more reside (T13) fully restored the response to carvacrol (Fig 3E). This result was consistent with results obtained with other activating MTs (not shown). The results further suggest the importance of three arginine residues (Arg344-346) in the response to MTs, as the deletion of even one of the three (T10) resulted in a channel that was indifferent to MT levels. We thus mutated the three arginine residues to either three alanine residues or three glutamine residues, to maintain the polarity of this region. This substitution was performed on both the full-length channel (3A and 3Q; Fig. 3D) and on the shortest channel that maintained a wild type-like response to carvacrol, T13 (T13-3A and T13-3Q). All mutants were functional and displayed potassium selective currents. Substitution of the three arginine residues reduced activation by carvacrol two-to three-fold, as compared with the wild type channel (Fig. 3F). Moreover, replacement of the arginine residues within the T13 deletion mutant almost completely abolished its responsiveness to carvacrol (Fig. 3F).

### 3.6. Defining the region of the C-terminal domain that regulates activation by known regulatory variables

As mentioned previously, the C-terminal domain of K_2P_2.1 channels was shown to regulate channel activity (Bagriantsev et al., 2012; Bockenhauer et al., 2001; Chemin et al., 2005; Gonzalez et al., 2015; Honore et al., 2002; Maingret et al., 2000; Maingret et al., 1999; Murbartian et al., 2005; Patel et al., 1998; Sandoz et al., 2012; Treptow and Klein, 2010). To further pinpoint the region within the C-terminal domain that governs the response to selected regulators, we tested the influence of various C-terminal domain deletions on regulation by arachidonic acid, temperature and holding potential changes. Accordingly, our array of truncated channels was tested for the effects of a 4 minute-long perfusion with 100 µM arachidonic acid, a temperature increase from 20°C to 35°C and a change in the holding potential from −20 to −80 mV (Cohen and Zilberberg, 2006; Segal-Hayoun et al., 2010). A full response was observed upon arachidonic acid treatment even with truncation T4 (Fig. 4A). The response to the temperature change was restored in channel variant T7 (Fig. 4B). K_2P_2.1 was previously shown to close when the membrane was held at −20 mV for prolonged times, whereas the channel was opened when the holding potential was shifted to −80 mV (Cohen and Zilberberg, 2006). Here, recovery of the response to changes in holding potential was achieved only in truncation T14 (Fig. 4C) and above. Substituting the three arginine residues important for responding to MTs to glutamines or alanines as before (Fig. 4D) abolished the response to changes in holding potential.

**Fig. 4.**
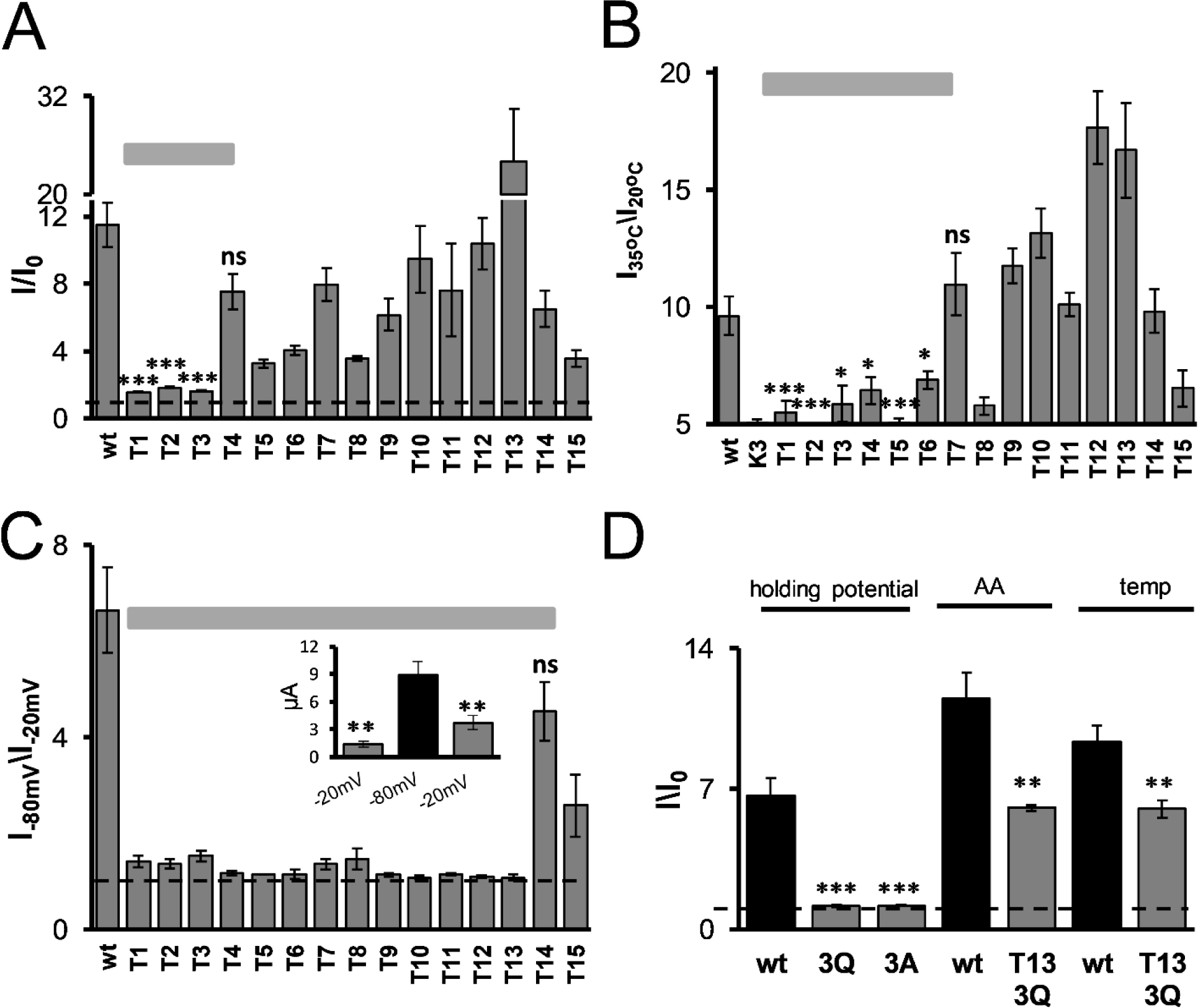
Delineating the minimal C-terminal segment essential for the response to regulatory variables. K_2P_2.1 channel responses to arachidonic acid, temperature changes, and membrane holding potential were tested as in Fig. 1A (n=5-10, mean±SEM). The dashed lines represent no change from the initial current. The horizontal bars indicate the minimal C-terminal domain that is essential for the response to the tested variable. (A) The response to arachidonic acid (100 µM, 4 minutes) is partially restored in the T4 variant (K_2P_2.1 1-337). (B) Response to temperature changes (20 to 35o C, 1.5 minutes) is restored in the T7 variant (K_2P_2.1 1-340). As a baseline, we used the current increase in a non-temperature sensitive channel, K_2P_3.1 (K3, 4.75-fold increase). (C) Current increase due to a change in the holding potential from −20 to −80mV (Cohen and Zilberberg, 2006; Segal-Hayoun et al., 2010) is measured only in the truncation variant T14 (K_2P_2.1 1-358) and above. Inset: average currents of wt K_2P_2.1 channels held at −20mV, −80mV and then again at −20mV. (D). Substitution of arginine residues at positions 344-346 by glutamines (3Q) or alanines (3A) completely abolished the response to holding potential changes, yet only had a minor effect on the responses to arachidonic acid and temperature change.

## 4. Discussion

In this study, we used an exogenous expression system to measure the impact of MTs on human K_2P_2.1 channel activity. We report, for the first time, that the activity of a K_2P_ channel is largely affected by MTs. We, moreover, described the physical and chemical requirements of a monoterpene that are essential for its ability to affect K_2P_2.1 channel activity and identified a distinct regulatory domain within the carboxyl-terminal domain of the channel that dictates its sensitivity to MTs.

A 40 percent increase in K_2P_2.1 channel current was observed upon treatment with 50 µM carvacrol, while an increase of 450% was detected in the presence of 900 µM carvacrol (EC_50_=200µM). Other MTs were found to affect various ion channels at comparable concentrations (Bavi et al., 2016; Brohawn et al., 2014; Cabanos et al., 2017; Joca et al., 2012; Johnson et al., 2012; Lauritzen et al., 2005; Li Fraine et al., 2017; Pham et al., 2015). It is thus plausible that the physiological effects of MTs are mediated, in part, by their activity on the K_2P_2.1 channel. For example, the known neuroprotective properties of carvacrol might be partially due to its effects on the K_2P_2.1 channel, a known target of neuroprotective agents (Duprat et al., 2000). In animal models, high micromolar concentrations of MTs were recorded after feeding with relatively large amounts of MTs or after injection of MTs intravenously (Ocel’ova et al., 2016; Pavan et al., 2018). It is conceivable that local high concentrations of MTs can be achieved using a topical application and high concentration of MTs in the plasma or certain tissues using an extensive oral administration, emulsification or β-cyclodextrins (β-CD) complexation (LeBlanc et al., 2008).

Our results also suggest a change in the selectivity of the channel as well as in its voltage dependence due to carvacrol application (Fig. 1E-F). These results are consistent with studies demonstrating that changes in K_2P_2.1 open probability affect both selectivity and voltage dependence (Bockenhauer et al., 2001; Cohen et al., 2008; Lopes et al., 2005).

Whether small amphiphilic molecules exert their activity on membrane proteins by direct binding (Nury et al., 2011; Ogawa et al., 2009) or by altering membrane properties which, in turn, enable conformational changes of the protein (Epand et al., 2015; Lee, 2011; Sacchi et al., 2015) remains an ongoing debate. We believe that the latter applies to the observed augmentation of K_2P_2.1 channel activity by terpenes and that modulation of channel activity is due to changes in the supramolecular organization of the membrane environment because a) a relatively high concentration of terpenes is needed to observe the effect on channel activity, b) many membrane proteins have been reported to be affected by terpenes at the same concentration range (Bavi et al., 2016; Brohawn et al., 2014; Cabanos et al., 2017; Joca et al., 2012; Johnson et al., 2012; Lauritzen et al., 2005; Li Fraine et al., 2017; Pham et al., 2015) and c) several terpenes with various structures had a similar effect on the channel. Reciprocal relationships between embedded proteins and the surrounding membrane arise from both specific and non-specific interactions. Membrane phospholipids might specifically interact with proteins or can serve as precursors for molecules that participate in protein regulation, such as arachidonic acid or PIP_2_ (Lundbaek et al., 2010). Transmembrane proteins might affect the membrane non-specifically through hydrophobic-matching and adjust its thickness (Harroun et al., 1999). At the same time, properties of the membrane, such as thickness, curvature, packing, lateral pressure and charge, influence the protein in a non-specific manner (Seeger et al., 2010). Those mechanisms were demonstrated for the KcSA channel, whose activity was found to be strongly related to the thermodynamic state of the membrane (Alessandrini and Facci, 2011), for the GABA_A_ receptor, where application of two neurosteroid isomers had opposite effects on bilayer properties (Sacchi et al., 2015), and for gramicidin A and voltage-gated sodium channels, that were affected by various structurally different amphiphilic drugs (Lundbaek et al., 2010).

Bilayer perturbation and reversible interactions of amphiphilic molecules, like terpenes, with the membrane, could decrease bilayer elasticity. Due to their “cone”-like shape, micelle-forming amphiphiles, such as thymol (Turina et al., 2006), could also elevate the degree of membrane curvature. It has been shown that major lipid properties, such as membrane fluidity, and other physiochemical properties of membranes are modified by MTs (Oz et al., 2015; Reiner et al., 2009; Sanchez et al., 2004; Turina et al., 2006; Zunino et al., 2011). These changes in lipid bilayer structure can affect the energetic requirements for gating-related conformational changes, as demonstrated for the nicotinic acetylcholine receptor (Barrantes et al., 2010; Fantini and Barrantes, 2009). Thus, terpenes are expected to non-specifically modulate the activity of membrane proteins. Here, we showed that MTs, which are amphipathic in nature, activate K_2P_2.1 channels and that small change in MTs structure that might affect their ability to perturb the membrane, caused dramatic differences in the ability to exert these effects. MTs can be either cyclic (aromatic or not) or linear, differing from one another by their hydrophobicity (expressed here as octanol-water partition coefficient (logP) prediction -XLogP3) and their polar area. By determining the impact of several terpenes with various properties on channel activity, we found that the degree of hydrophobicity and polarity, as well as the structure of the monoterpene, were all crucial to their effect on the channel. For maximal impact on K_2P_2.1 channel activity, terpenes should be moderately hydrophobic (XLogP3 ~ 3, as is the case for carvacrol, thymol, and 4-IPP) and to be able to penetrate yet not become embedded in the bilayer. Similarly, eliminating the polar area (p-cymene) led to a substantial decrease in the ability of a monoterpene to open the channel. In addition, a phenol moiety was necessary to obtain high channel-stimulating activity. Accordingly, menthol was much less effective than was thymol. Additionally, other traits of the terpene should be considered as they are also expected to change membrane phospholipids parameters, such as area per lipid, thickness, order parameters and the tilt of the *sn*-1 and *sn*-2 chains (Witzke et al., 2010). This is consistent with the finding that small and rigid amphipathic molecules are likely to penetrate and increase membrane lipid acyl chain hydrocarbon group order near the interface (Pham et al., 2015) and that when integrated into membranes, hydroxyl-containing MTs are located within the vicinity of the polar glycerol/phosphate backbone of the bilayer (Witzke et al., 2010). Furthermore, it was demonstrated that localization of alcohol-based MTs near the polar membrane area increased membrane curvature, whereas partitioning hydrocarbons into the more hydrophobic membrane core can cause membrane expansion and stiffening (Zunino et al., 2011).

To learn whether MTs entry to the extracellular facing membrane affects K_2P_2.1 channels through their membrane-embedded domain or by changing the interaction of their regulatory C-terminal domain with the membrane, we first tested the effect of complete removal of the cytoplasmically-oriented carboxyl-terminal domain of the channel. Since the truncated channels were not affected by added terpenes, we assume that the interaction between the inter-membrane domain of the channel and the bilayer is not sufficient to communicate membranal perturbations into protein-gating movements. Similarly, the carboxyl-terminal was necessary for stretch activation of the channel (Maingret et al., 1999). Thus, for the gating mechanism of K_2P_2.1 channels to follow the “force-from lipids” paradigm (Bavi et al., 2016), the proximal region of the carboxyl-terminal domain is needed. Mechanosensitivity of the channel was found to be mediated directly by the lipid membrane (Brohawn et al., 2014), even though, inhibition of channel activity by the cytoskeleton was reported as well (Lauritzen et al., 2005; Li Fraine et al., 2017). To locate the domain that is fundamental to monoterpene sensitivity, we tested a series of mutated channels with sequential truncations of the carboxyl-terminal domain (Fig. 3). We found that the proximal 33 residues of this domain are sufficient for activation of the channel by MTs (T12, Fig.3E).

The carboxyl-terminal domain described here is longer than the domain previously described as being essential for regulation of the K_2P_2.1 channel, with Glu321 being found to facilitate internal pH sensing (Honore et al., 2002), a patch of positively charged residues, Arg312, Lys316, Lys317, and Lys319 have been implicated in PIP_2_ modulation (Cabanos et al., 2017; Chemin et al., 2005) and a stretch of amino acids up to Thr337 has been reported as necessary for activation by arachidonic acid (Patel et al., 1998). We thus determined the minimal region of the carboxyl-terminal domain important for activation by temperature, arachidonic acid and membrane potential (Fig. 4). We were able to locate two distinct regions, a more proximal domain (up to residue Lys341) that is necessary for activation by arachidonic acid anionic phospholipids and temperature and one more distal (up to residue Ala358) that is essential for activation by MTs and holding potential changes. A cluster of three arginine residues (residues 344-6) was found to be significant not only for MTs-mediated activation of the channel but was also essential for sensing membrane potential changes (Figs. 3 and 4). In other membrane proteins, “poly-basic clusters” were reported to induce binding to PIP_2_ (Johnson et al., 2012; Kaur et al., 2015). As the sensitivity of the K_2P_2.1 channel to membrane potential changes was reported to be due to changes in PIP_2_ levels, following fluctuations in the activity of phospholipase C (Segal-Hayoun et al., 2010), our findings could imply that the three arginine residues constitute a second PIP_2_-binding site. Indeed, the importance of this domain for the basal activity of K_2P_2.1 and K_2P_10.1, but not for that of K_2P_4.1 lacking this poly-arginine motif, was demonstrated of late (Soussia et al., 2018). Likewise, Woo et al. (Woo et al., 2018) also concluded that this motif is a high-affinity PIP_2_-binding site, representing a candidate region for stimulatory regulation by lower levels of PIP_2_. It should be noted that MTs and membrane-holding potential changes do not operate through identical mechanisms as the phosphorylation of Ser348 affects channel responsiveness to the holding potential (Segal-Hayoun et al., 2010) but not to MTs (Fig. 4). This point should be further investigated. We thus believe that by embedding into the membrane and changing its properties, MTs affects the interaction of the C-terminal of the channel with the intracellular side of the membrane, resulting in current augmentation.

Integration of our results with those of previous studies, summarized by Woo et al. (Woo et al., 2018), led us to suggest a comprehensive model explaining the contribution of the interaction between the carboxyl-terminal domain of the K_2P_2.1 channel with the membrane to channel regulation and gating (Fig. 5). According to this integrated model, the basal activity and regulation of the channel are dictated by the binding of two carboxyl-terminal domains to the membrane, one proximal and one distal to TM4. The former presents a low-affinity PIP_2_-binding domain, while the latter presents a high-affinity PIP_2_-binding domain within an arginine cluster. When both domains are not in contact with the membrane due to either removal of the majority of the carboxyl-terminal (Figs. 3 and 4) or the addition of poly-L-lysine (PLL, Fig. 5E), channel activity is low and cannot be regulated. The activity also cannot be regulated when the channel is “locked” in a constantly active state. This occurs when the pH sensor, Glu321, is neutralized, regardless of the location of the distal domain, (Fig. 5D) or when PIP_2_ levels in the membrane decrease, which disables binding of the proximal domain to the membrane (Fig. 5B). The proximal domain is essential for complete regulation by monoterpenes and membrane voltage changes, as elimination of this domain (Figs. 3E and 4C) or replacement of its poly-arginine cluster (Figs. 3F and 4D) attenuates the ability of the channel to be regulated by both factors (Figs. 5A and 5C).

**Fig. 5.**
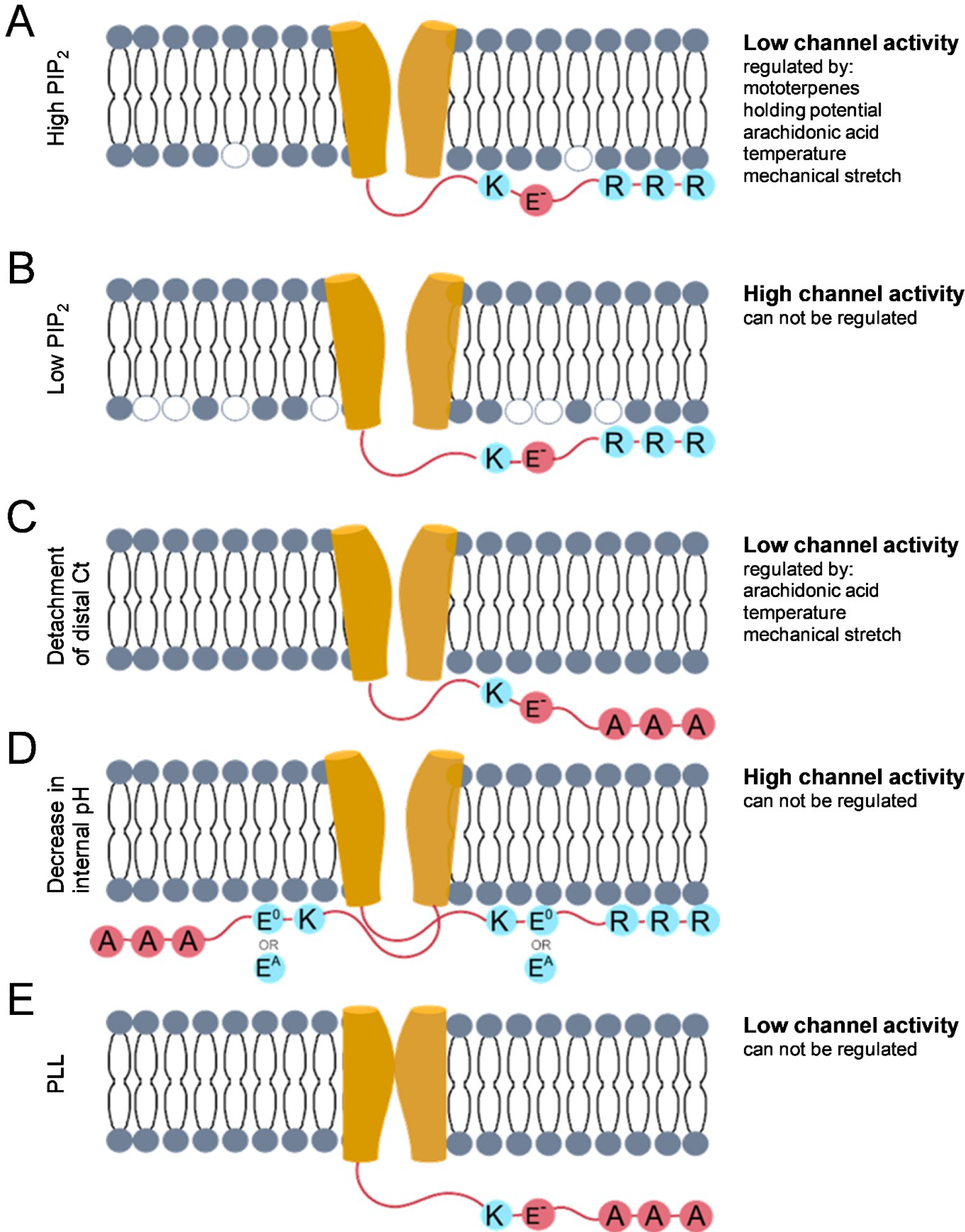
A schematic model of the interplay between the different C-terminal domains and the sensitivity of K_2P_2.1 channels to various stimuli. According to the suggested models for K_2P_2.1 (Chemin et al., 2005; Honore, 2007) and K_2P_10.1 (Woo et al., 2018; Woo et al., 2016) channel regulation, the activity level of the channel is tightly related to the interaction of two intracellular C-terminal domains (a proximal one and a distal one) with the membrane. The proximal domain was proposed to contain a low-affinity site for PIP_2_ binding, while the distal domain was proposed to present a high-affinity site for PIP_2_ binding (Woo et al., 2018). The presented model is based on previous models proposed for both channels, as well as on the results described in the present study. (A) Under standard conditions (relatively high PIP_2_ levels), the distal domain is bound to the membrane, while the proximal domain is partially bound. Under these conditions, channel basal activity is low yet can be enhanced by the addition of MTs or arachidonic acid, as well as by increased temperature, mechanical force and changes in the membrane-holding potential. (B) Reduction in PIP_2_ levels promotes detachment of only the proximal domain from the membrane, resulting in a constantly active channel. (C) Removal of the distal domain results in low channel activity and completely abolishes the response to MTs and changes in the membrane-holding potential. Neutralization of a poly-arginine motif within the distal domain abolishes channel regulation via changes in the membrane-holding potential and reduces the effect of MTs. Both modifications do not alter responses to other regulators. (D) Neutralization of an internal pH sensor (Glu321) results in a constantly active channel, regardless of the position of the distal domain. (E) When both domains are detached from the membrane, with the addition of poly-L-lysines (PLL), the channel is inactive and cannot be opened.

Using MTs, we were able to detect a novel regulatory domain in the carboxyl-terminal domain of the K_2P_2.1 channel. Moreover, the growing number of studies on the use of MTs for the treatment of neuropathy (Abuhamdah et al., 2015; Porres-Martinez et al., 2016), as well as for treating diabetes (Habtemariam, 2017), points to the need for better understanding of the effect of MTs on cell components, such as K_2P_ channels, which are involved in these conditions.

## Acknowledgments

The authors thank Profs. Itzhak Fishov and Oded Farago for their advice and suggestions.

